# New to Ehretiaceae: *Keraunea*. Re-evaluation of a genus of climbers from Brazil

**DOI:** 10.1101/2023.02.09.527833

**Authors:** Martin Cheek, Julio A. Lombardi, Ana Rita G. Simões, Alexandre R. Zuntini

## Abstract

We definitively place *Keraunea*, a genus of showy forest climbers from remnants of the Mata Atlântica of Brazil, in Ehretiaceae. Previously *Keraunea* had been ascribed to Convolvulaceae based on morphology, or divided between Malpighiaceae and Ehretiaceae based on molecular analyses (polyphyletic). *Keraunea* is morphologically anomalous in the Ehretiaceae, having fruits which are held in the centre of a large wing-like bract by adnation of the pedicel, and due the stem-twining habit of some species. However, morphologically *Keraunea* shares two apomorphies with monotypic *Cortesia* Cav., halophytic shrubs of Argentina: 1) “two parted endocarps” (the fruit contains two endocarps each with two 1-seeded cells, while four 1-seeded endocarps are plesiomorphic in Ehretiaceae) and 2) a large bract that subtends the flower (absence of bracts is plesiomorphic in Ehretiaceae). A combined tree using four markers (ITS, *trn*L-F, *rbc*L and *mat*K) places the three species of *Keraunea* sampled unambiguously in a clade with *Ehretia* + *Cortesia* + *Halgania* and monophyly for *Keraunea* is shown with high support. In an ITS tree *Keraunea* is sister to *Cortesia* with low support.

We expand the generic description of *Keraunea* including the first account of the internal fruit structures and the seed, and present new data on the vegetative architecture including variation within the genus, some species being stem twiners while one species at least, is described as a scandent shrub. *Keraunea lombardiana*, previously included within *K. brasiliensis*, is formally described from Minas Gerais and Bahia as the third species of the genus and both these species are described. In all five species are recognised but two, known to us only from digital images, are not formally described because sufficiently detailed descriptions cannot be made. All five species are mapped, and provisional conservation assessments are recorded, of either Endangered or Critically Endangered. The state of Bahia, with three species, has the highest species diversity, mainly in dry forest. Three species appear confined to moist coastal forest, extending southwards from Bahia to the state of Rio de Janeiro.

*Keraunea* remains incompletely known. Not one of the species has both anthetic flowers and fruits described. Studies on pollen, germination, anatomy, embryology and phytochemisty are entirely lacking. Field observations of pollination, seed dispersal and phenology are also required. However, the most urgent requirement is undoubtedly a formal taxonomic revision based on a full herbarium search and targeted fieldwork, with full IUCN 2012 extinction risk assessments for each taxon. This is urgent because it seems that all the taxa that we present in this paper (and more that can be expected to be found) appear to be highly range restricted and generally not to occur in protected areas, and so appear to be highly threatened.

## Introduction

*Keraunea* Cheek & Sim.-Bianch. based on *K. brasiliensis* Cheek & Sim.-Bianch. was published for a new genus and species of Brazilian climber with seed dispersal aided by a wing-like bract to which the fruit is attached centrally by adnation of the pedicel (Cheek & Simão-Bianchini 2013). The type specimen arose from botanical survey work for a checklist of the Catolés area in the Chapada Diamantina, Bahia (Zappi *et al*. 2006).

The genus was placed in Convolvulaceae because the unusual fruit arrangement is identical to that of *Neuropeltis* Wall. of the Old World (Africa and Asia). Placement in the Convolvulaceae was supported by the superior ovary with 2 carpels, a bifid stigma, gamopetalous corolla, and epipetalous stamens, climbing habit, with alternate, exstipulate, pinnately nerved, simple, entire leaves. Placement within the existing infra-familial classification of Convolvulaceae was problematic. The possibility of convergent evolution was considered, but was set aside (Cheek & Simão-Bianchini 2013). A second species was soon published, *Keraunea capixaba* Lombardi (Lombardi 2014) (Fig. 1). *Keraunea* was accepted and widely cited in botanical literature (e.g. Costa *et al*. 2015; Simão-Bianchini *et al*. 2015; Marinho *et al*. 2016; Simão-Bianchini (2020); Alencar *et al*. 2022). However, Simões *et al*. (2022), as part of a programme using genomic data to complete the tree of life at generic level (the Plant and Fungal Tree of Life PAFToL e.g. Baker *et al*. 2021) using a sample from *Keraunea capixaba*, suggested that the genus should be placed outside the Convolvulaceae, but did not suggest an alternative familial placement. Independently, researchers using sanger sequencing methods also excluded *Keraunea* from Convolvulaceae (Muñoz-Rodriguez *et al*. 2022). They concluded that *Keraunea brasiliensis* should be placed in Malpighiaceae, while *Keraunea capixaba* should be placed in Ehretiaceae. Thus, they concluded that the genus was not monophyletic, given that, in their analysis, the type specimen of *Keraunea brasiliensis, Passos* 5263 was most likely a *Mascagnia* (DC.) Bertero and would justify transferring *K. brasiliensis* to this genus (Muñoz-Rodriguez *et al*. 2022). However, this attribution to Malpighiaceae of *Keraunea brasiliensis* was recently shown to be due to an uncritical sampling error (de Almeida *et al*. 2023).

**Fig. 1.**
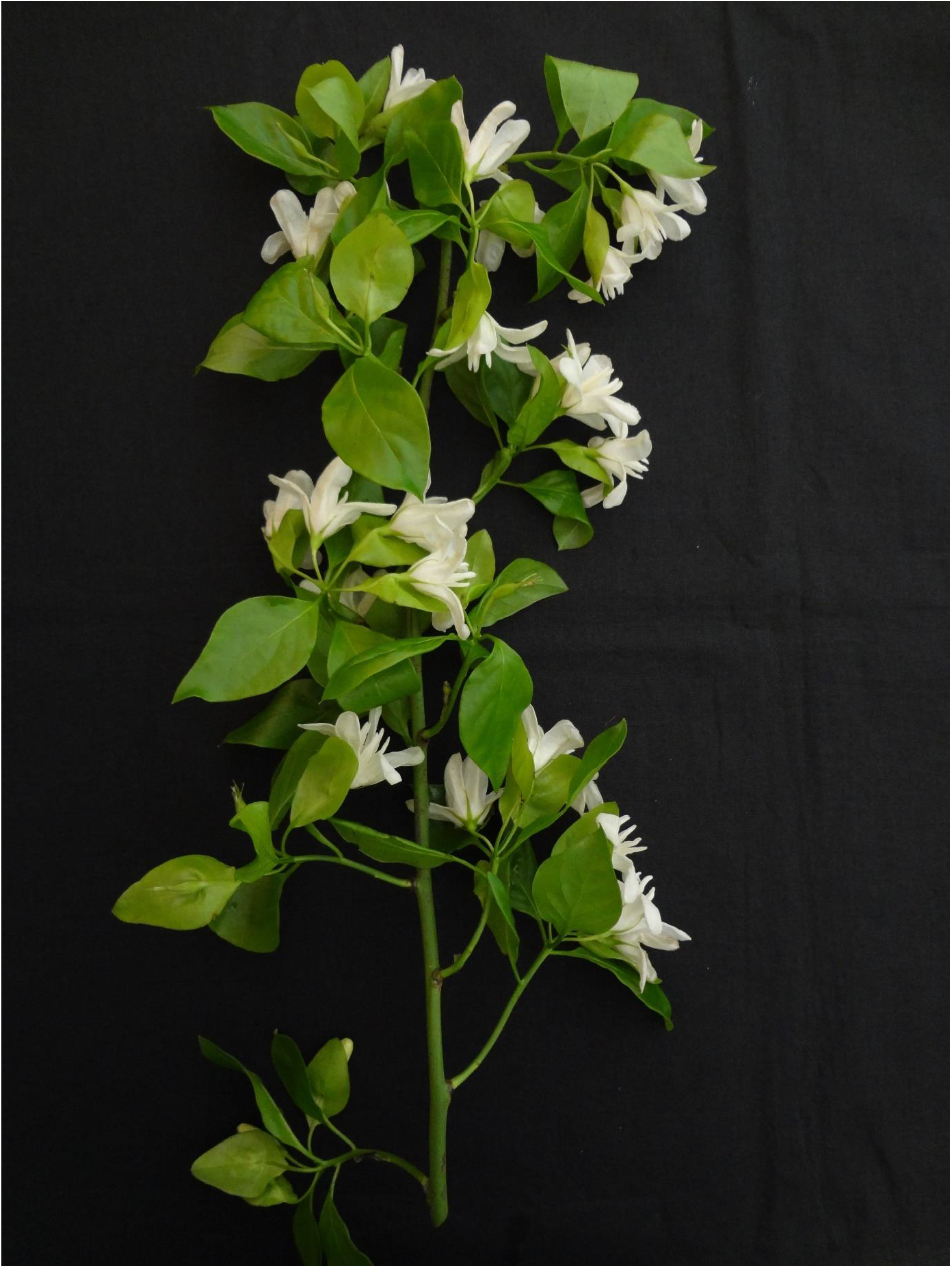
*Keraunea capixaba*. Primary (extension) shoot with numerous flowering spur-shoots. Note that some flowers lack bracts. From *Siqueira* 891 (type). Photo by Geovane Siqueira each other, and sister to *Ehretia*, with high support. However, neither *Cortesia* nor *Halgania* have been sequenced for *mat*K.

In contrast, sanger sequencing of two other specimens of *Keraunea* (*Lombardi* 1819, paratype of *K. brasiliensis, Siqueira* 893, paratype of *K. capixaba*) placed them in the Ehretiaceae (Boraginales) (Muñoz-Rodriguez *et al*. 2022). In a densely sampled *nr*ITS phylogeny of Ehretiaceae, *Lombardi* 1819 is sister to monophyletic *Cortesia* Cav., forming a clade that is sister to the Australian genus *Halgania* Gaudich. with moderately high support. *Cortesia* and *Halgania* form a clade that is sister to *Ehretia* P.Br. In the *mat*K phylogeny, the two *Keraunea* specimens (*Lombardi* 1819 and *Siqueira* 893) are most closely related to

Ehretiaceae has been delimited in different ways since it was resurrected from synonymy within a widely defined Boraginaceae. In this paper we follow the delimitation of the familial classification of the Boraginales by the 20 members of the Boraginales Working Group which recognises 11 families (Luebert *et al*. 2016) rather than that of Gottschling *et al*. (2016) in which Cordiaceae and Hoplestigmataceae are included in Ehretiaceae.

Here, we review and extend the morphological data available for *Keraunea*, examining the fruits and seed in detail for the first time and presenting an expanded description of the genus. We test morphological and molecular data, addressing: 1) the placement of *Keraunea* in Ehretiaceae; 2) its relationships within the family; 3) the monophyly of the genus. We then formally transfer *Keraunea* to Ehretiaceae, and update knowledge of the genus, including the formal description of a new, third species, with notes on two further, imperfectly known species.

## Materials & Methods

Herbarium material was examined with a Leica Wild M8 dissecting binocular microscope fitted with an eyepiece graticule measuring in units of 0.025 mm at maximum magnification. The drawings were made with the same equipment with a Leica 308700 camera lucida attachment. Fruits were soaked overnight in hot water before dissection. Photographs of fruit structures were taken with a digital camera coupled to a Leica S9i stereomicroscope. Specimen data and images were studied on Lista do Brasil-online (continuously updated) and through gbif.org. Fifty-one occurrence records representing 19 specimen numbers were discovered and downloaded as the following dataset from GBIF.org (24 January 2023) GBIF Occurrence Download https://doi.org/10.15468/dl.6bpx4k. Herbarium codes for specimens as indicated in the results section follow Index Herbariorum (Thiers continuously updated). Specimens cited have been seen unless indicated “n.v.”. Names of species and authors follow the International Plant Names Index (IPNI continuously updated). Nomenclature follows Turland *et al*. (2018). Technical terms follow Beentje & Cheek (2003). The conservation assessments follow the IUCN (2012) categories and criteria. A distribution map of all the known species of *Keraunea*, including the newly proposed ones, was prepared based on geographic coordinates available from herbarium specimens. When coordinates were not available, these were visually estimated in Google Maps using the gazetteer descriptions provided in the collector’s notes (Appendix I); an ecological layer representing the extent of Mata Atlântica was projected on the species distribution map, based on data from Oliveira Filho (2014).

To evaluate the phylogenetic placement of *Keraunea*, new DNA sequences were produced from three specimens at Kew: an isotype of *K. brasiliensis* (*Passos* PCD 5263), a paratype of *K. capixaba* (*Siqueira* 893), and a third specimen (*Lombardi* 1819), here described as a new species, *K. lombardiana*. Leaf fragments were carefully sourced from specimen envelopes. Total DNA was extracted using CTAB (Doyle & Doyle 1987) and purified using NucleoSpin® Purification Kit (Macherey-Nagel, Düren, Germany). DNA extractions were profiled in Agarose gel 1% and using Nanodrop. Four markers were amplified: ITS, *trn*L-F, *rbc*L and *mat*K, with primers and conditions specified in (Supplementary Material). All PCR reactions were performed in 50µl volume containing: 25µl of Taq PCR Master Mix (QIAGEN, Germany), 2.5µl of BSA (0.2 μg/μL), 2.5µl of DMSO (5%), and 1µl of each primer (0.2 μM). Sequences were assembled in Geneious 8.1.9 (www.geneious.com) and then blasted against NCBI database, using MEGA BLAST (Camacho et al, 2009); to avoid circularity in the results, we excluded the sequences of *Keraunea* from the reference. Assembled sequences were annotated using *Ehretia* sequences as reference and deposited in NCBI.

To produce the matrices for the four markers, all sequences of Ehretiaceae available in the NCBI were pre-selected and only one accession per species was kept, matching as much as possible the same specimen across the four markers. Outgroups were selected based on Gottschling *et al*. (2014), with sequences for Cordiaceae and Heliotropiaceae also obtained from NCBI. The list of accessions used is available (Supplementary Material). For each marker, an alignment was produced in Muscle (Edgar 2004) using standard parameters, and manually trimmed in Geneious. The four matrices were analysed together in IQ-Tree 2.2.0 (Nguyen *et al*. 2015), using ultrafast bootstrap (BS) as branch support (Hoang *et al*. 2018). A second analysis with the ITS matrix alone was also produced (Supplementary Material).

## Results

### Molecular phylogenetic analyses

Twelve novel sequences were produced, ranging from 584bp to 890bp, and all of these had Erhetiaceae as top hit; for matK, *Cordia* and *Ehretia* had identical maximum scores. Sequences of matK and rbcL had the highest similarity to the reference, respectively 99% and 98%, followed by trnL-F (90-94%) and ITS (84-86%). The list of 10 best hits per sequence are available in the Supplementary Materials.

Our phylogeny reconstructs Ehretiaceae as a highly supported clade (100% Bootstrap Support or BS), sister to Cordiaceae (Fig. 2). *Keraunea* is demonstrated to belong in Ehretiaceae, and resolved as sister to *Halgania*, with 90% BS, and the two genera are resolved as sister to *Cortesia*; however, this relationship is not as strongly supported (81% BS). In the ITS tree (Suppl. Material), *Keraunea* emerges as sister to *Cortesia* (78% BS), both sister to *Halgania* (81% BS).

**Fig. 2.**
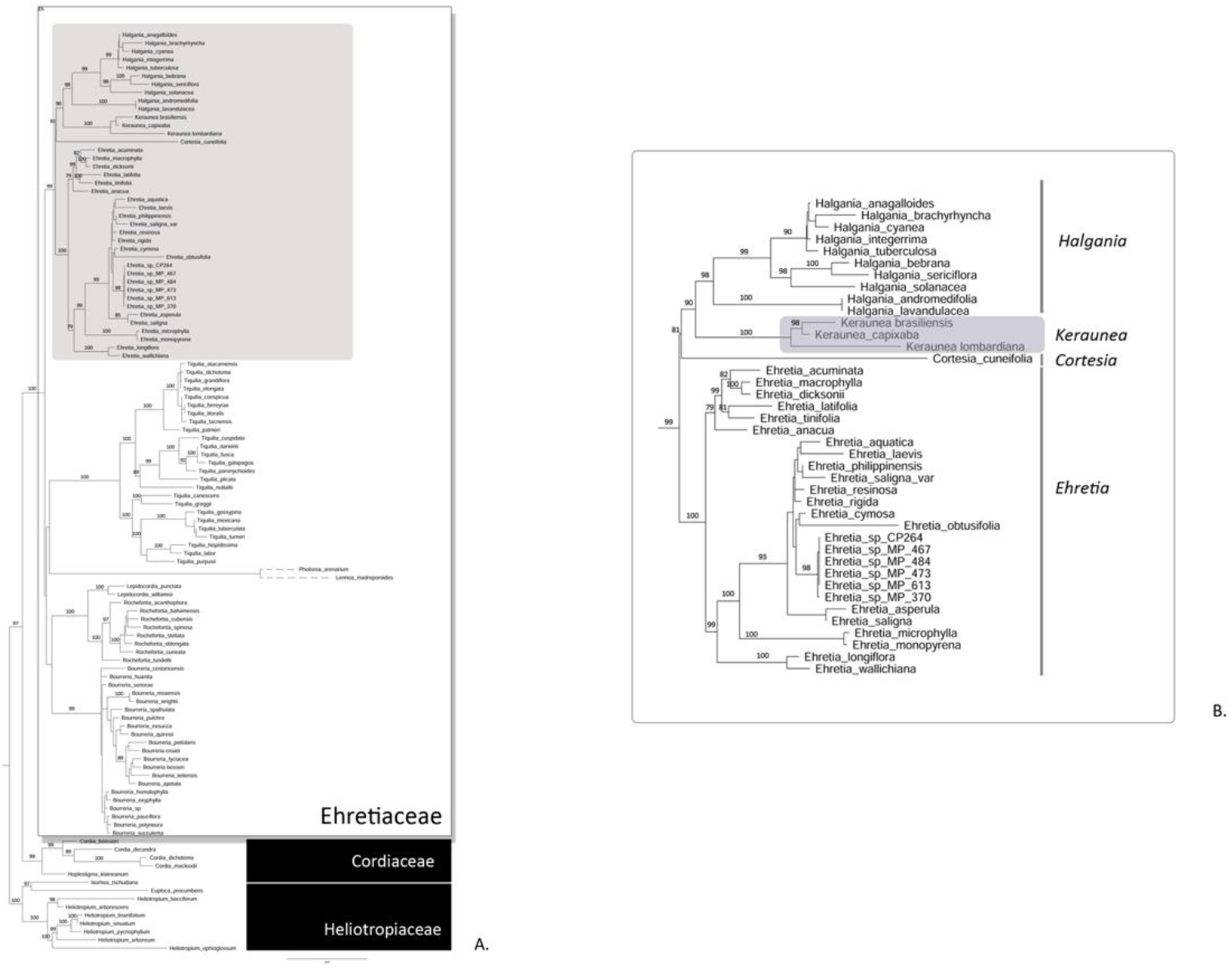
Phylogenetic hypothesis of Ehretiaceae, now including *Keraunea*. Maximum likelihood phylogram, based on the concatenated matrix composed of ITS, trnL-F, rbcL, and matK, with branch support inferred using ultrafast bootstrap (as percentage).

The monophyly of the genus is also strongly supported (100% BS), therefore dismissing previous hypotheses that it may be polyphyletic. The molecular analyses also show *K. lombardiana* as sister to the clade that includes *K. brasiliensis* and *K. capixaba*, further supporting its distinctiveness from *K. brasiliensis* and the value of recognising it as a new species to science.

### Taxonomy

***Keraunea*** Cheek & Sim.-Bianch.

#### Type species

*Keraunea brasiliensis* Cheek & Sim.-Bianch.

##### Hermaphrodite evergreen scandent shrubs or twining woody climbers

Primary axes (extension shoots) terete, fistular or solid, glabrous or hairy with more than one hair type; phyllotaxy spiralled, but subdistichous (approaching ½), unbranched (except in *K. brasiliensis*). Brachyblasts (spur-shoots) emerging from the primary axis from above the axillary buds, subtended by several bud-scales; spur shoots each with 2 – 4 dimorphic leaves and a terminal inflorescence. Leaves simple, phyllotaxy spiralled, petiolate, exstipulate, entire, pinnately nerved, lacking domatia, more or less brochidodromous; usually trimorphic, those of the primary axes different in size and shape to those of the spur shoots; the spur shoots with the proximal leaf usually smaller and differently shaped from the distal leaves. Petioles canaliculate, articulate at the junction with the stem, often slightly twisted. Indumentum varying between primary stem, brachyblasts, leaves and inflorescence to calyx (but nearly absent in *K. capixaba*) with up to two trichome types: 1) simple unicellular white or translucent hairs with an acute apex, erect or appressed, straight or curved or crisped, (uniquely in *K. lombardiana* stout, thickly cylindrical, sometimes from a raised, conical base and fracturing with age above the base, becoming scabrid); 2) uniseriate, erect, translucent, truncate hairs, the distal cell sometimes red. *Inflorescence* corymbose, subumbellate, 3(– 5)-flowered, terminal on short, leafy, spur-shoots, pedunculate, rhachis contracted. Pedicels articulated at base (*Keraunea lombardiana*), bract inserted at apex of pedicel, elliptic, base decurrent. Bracteoles absent or inconspicuous. Calyx tube campanulate, short, sepals imbricate, five, narrowly triangular to narrowly lanceolate, pinnately nerved, outer surface and margin with indumentum as leaf-blade, inner surface glabrous. Corolla (Fig. 3) white and sweetly scented at anthesis (where known), tube shorter than petals; petals convolute in bud, porrect, oblong-elliptic, apex rounded, sometimes with a few simple hairs near the apex, midline darker, glabrous; androecium epipetalous, stamens 5, inserted towards base of corolla tube, alternating with petals, glabrous, anthers mostly or fully exserted at anthesis; filaments cylindrical, anthers basifixed, cells introrse, sterile connective distal; disc intra-staminal, annular, sometimes inconspicuous; gynoecium superior, glabrous, ovary subglobose, style single, conduplicate, minutely bifurcate at apex, stigmas exserted, two, truncate or subcapitate; ovary 2-locular, locules biovulate, each becoming divided by a septum into two uniovulate cells, ovules anatropous, pendulous. *Infructescence* 1 – 3-fruited. Fruit usually inserted near the centre of the accrescent bract by adnation of the distal part of the pedicel; bract accrescent, papery, elliptic, wing-like; calyx persistent, green, not accrescent. Fruit green when ripe, drying brown, subglobose to broadly ellipsoid, glabrous, smooth or ridged-reticulate due to shrinkage onto the endocarp ornamentation when dried, style persistent; mesocarp spongiose with branched vascular bundles and little pulp; pyrenes two, plano-convex, appressed to each other by the flat ventral faces, dorsal surface convex and decorated with wings, ridges and/or spines, with or without a longitudinal groove dividing two collateral 1-seeded compartments. Seeds slightly curved, white, embryo about equal to endosperm, cotyledons flat, collateral, radical apical.

**Fig. 3.**
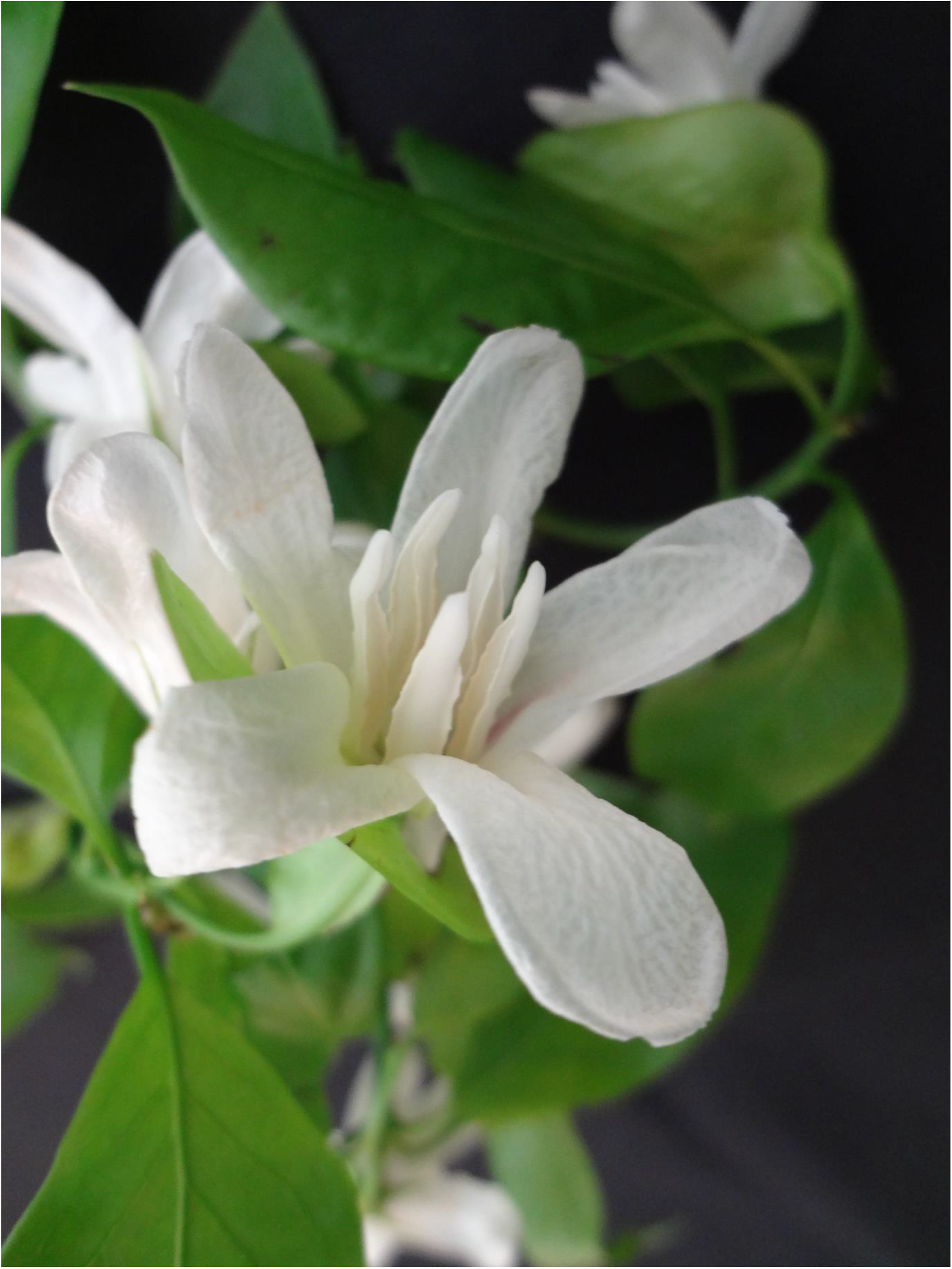
*Keraunea capixaba*. Close-up of flower. From *Siqueira* 891 (type). Photo by Geovane Siqueira

### DISTRIBUTION. BRAZIL

The states of Minas Gerais, Bahia, Rio de Janeiro and Espírito Santo. The centre of species diversity on current evidence is the state of Bahia, with three of the five known species. The other three states have a single species each.

### HABITAT

Dry, seasonal or mesophilic forest, often disturbed, altitudinal range 100 – 600 m.

### CONSERVATION ASSESSMENT

All known species appear range-restricted and are likely threatened. *Keraunea capixaba* was evaluated as Endangered using the IUCN (2012) criteria by Lombardi (2014). The other three taxa all appear to have fewer locations, probably smaller range sizes, and also face threats so are also expected to be highly threatened with extinction.

### PHENOLOGY

Flowering September-November, fruiting October to June.

### ETYMOLOGY

Named from the Greek for lightning bolt, to convey the unexpected nature of the discovery of this new genus in Brazil (Cheek & Simão-Bianchini 2013).

### VERNACULAR NAMES & USES

Only one species is reported to have a local name (Lombardi 2014) and no uses are reported.

### NOTES

Lombardi (2014) reported that in many inflorescences of *Keraunea capixaba*, one of the flowers often lacks a bract (see Fig. 1). This can also be seen in the specimen of *K*. sp. A.

Fruit colour is reported as green in the field notes and the fruits understandably stated to be immature. However, rehydration and dissection of the largest fruits of two species showed that their endocarps were fully formed, and their seeds mature or near mature (see species accounts below). Therefore, it appears that mature fruits in the genus may be green.

The endocarps have been investigated in two species and show characters that appear to be of taxonomic value at specific level in size and sculpture (wings and spines), and the extent of separation of the two seed cells from each other (Fig. 4, 5 & 7I).

**Fig. 4.**
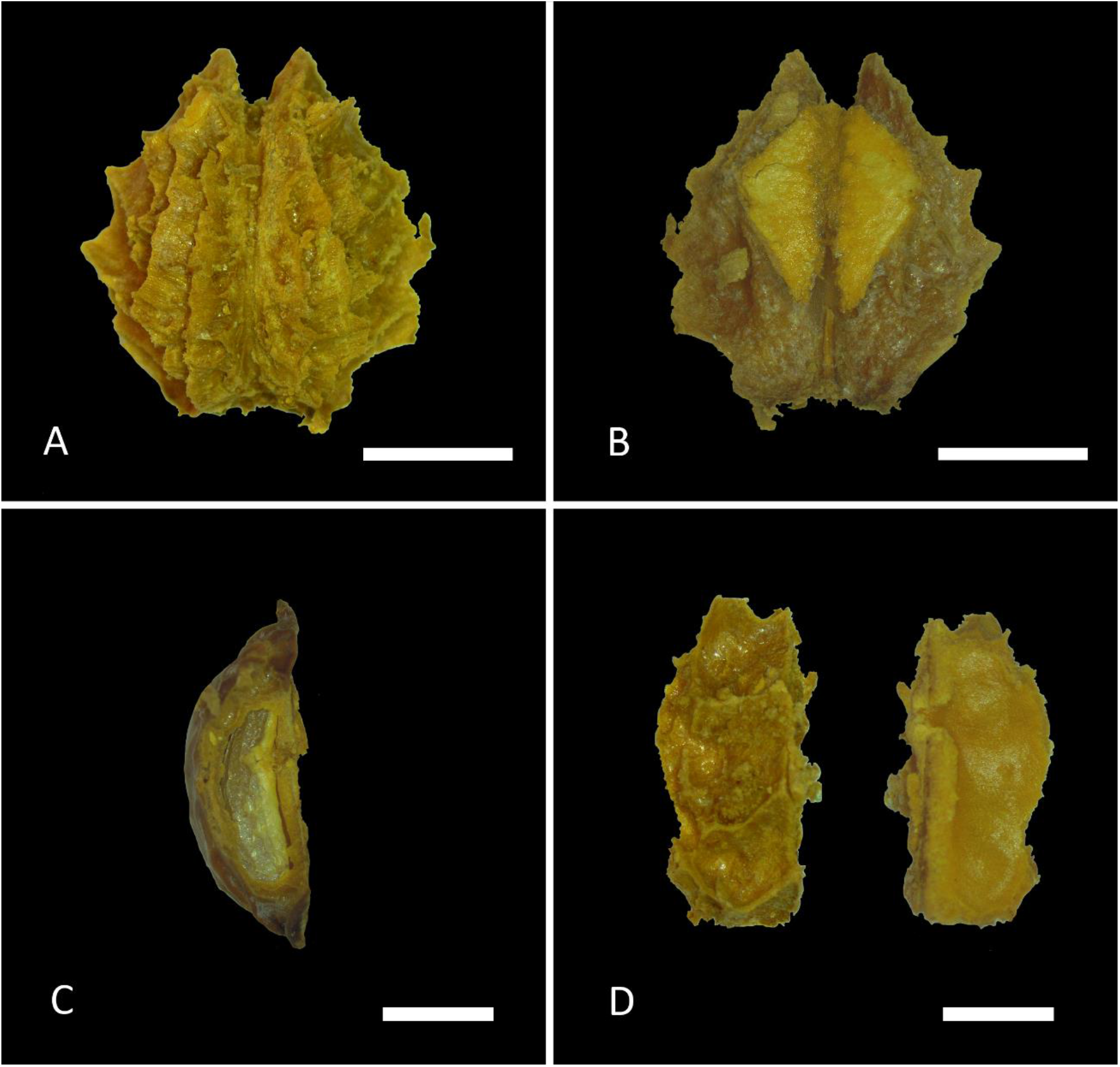
*Keraunea brasiliensis* “two-parted endocarps” (one of two from the fruit) A. dorsal face (convex) showing the central longitudinal groove, two apical and several lateral spines; B ventral surface (flat); C longitudinal section of one of the two seed compartments showing the slightly curved, white seed in side view; D side view of the endocarps. Scale bars = 2 mm (A&B) and 1 mm (C&D). All from *Passos et al*. 5263 (K). Photos by A.R.G. Simões

**Fig. 5.**
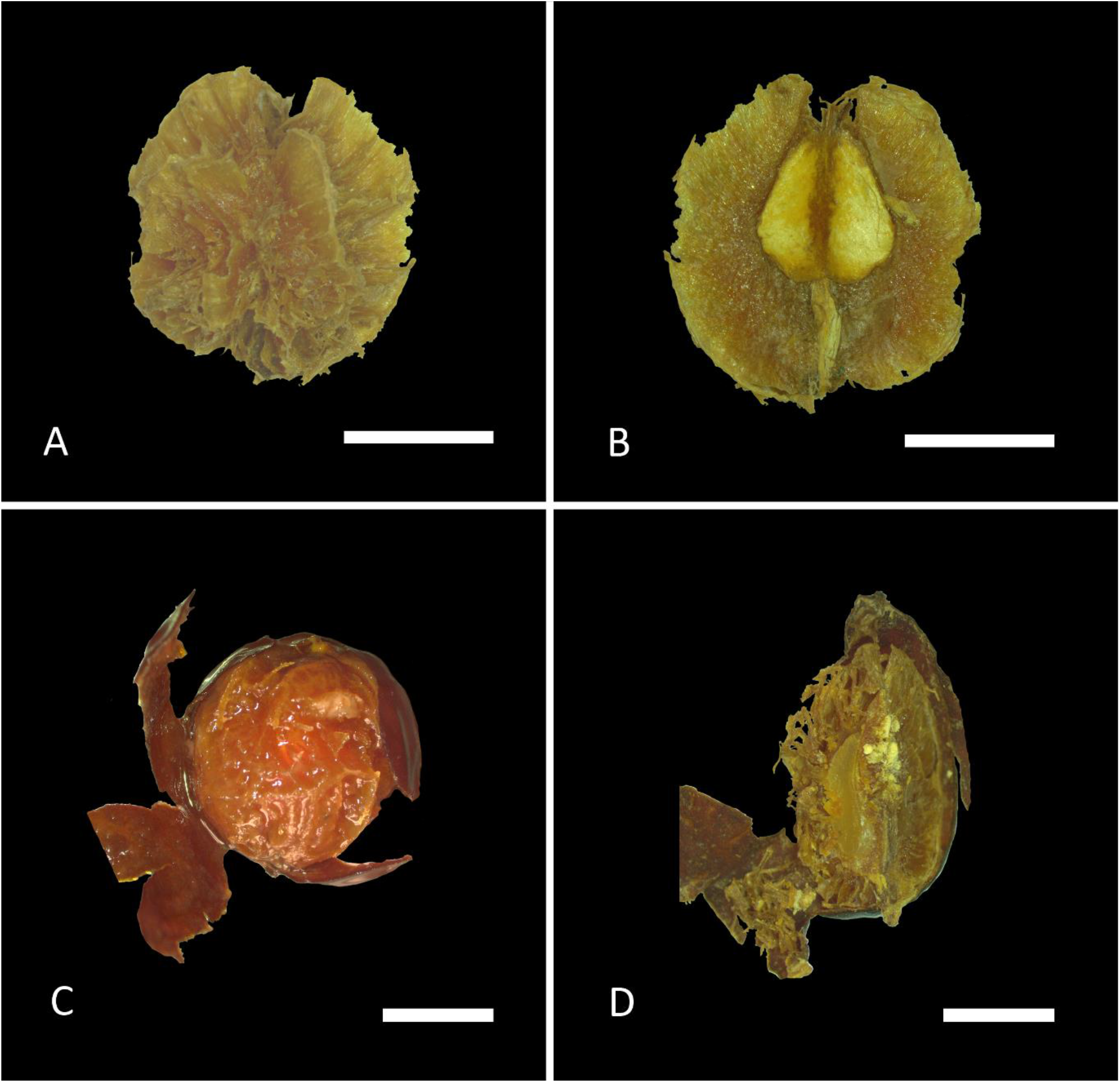
*Keraunea lombardiana* Fruit and endocarps. A. endocarp dorsal face (convex) showing the marginal wing, absence of spines and a central longitudinal groove; B endocarp ventral surface (flat); C fruit rehydrated with exocarp peeled to show the spongiose mesocarp; D fruit, sagittal section of showing the exocarp, spongiose mesocarp, and woody endocarp enclosing one of the seed compartments (seed, side view). Scale bars = 2 mm (A&B) and 1 mm (C&D). All from *Lombardi* 1819 (K). Photos by A.R.G. Simões

The soft mesocarp (of rehydrated fruits) under the thinly leathery epicarp is due to a mattress-like branched vascular system, with only a few pulpy cells. Presumably the fruit is dispersed when a gust of wind catches and transports the light wing-like bract to which the fruit is firmly attached, the base of the pedicel parting by abscission. When the fruit settles at its destination, it is likely that the mesocarp rots, releasing its two, 2-seeded endocarps, since the fruit lacks both attractive colour and food rewards for animal dispersers.

We speculate that the flowers might be pollinated by moths since where known they are both highly perfumed and bright white (Lombardi 2014), with a tube. However, field observations are needed to confirm this.

Where habitat quality is detailed in *Keraunea* specimen metadata, it is often reported that forest habitat is secondary or fragmentary, or that the specimen was collected from a forest edge. However, this may reflect that flowering plants of the genus, which appear to be canopy-flowering, are more easily detected and collected in such settings rather than being limited to them. A similar scenario has been reported in other rare, newly discovered canopy climbers (Cheek *et al*. 2022).

*Keraunea capixaba* roots easily from stem cuttings. Plants in cultivation have reached about 3 m tall after about 10 years, but have not yet flowered. The plants are evergreen and stems twine (Lombardi pers. obs.).

The species of *Keraunea* all appear to be restricted to the Mata Atlântica biome, of which <7% survives, and 80% of that exists in fragments of < 0.5 square kilometres (Hance & Butler continuously updated). The five species known to date occur in different forest types: atlantic moist forests (*K. capixaba, K. sp*.A and *K. sp*.B) and atlantic dry (deciduous) forests, (*K. brasiliensis* and *K. lombardiana*) (Map 1).

**Map 1.**
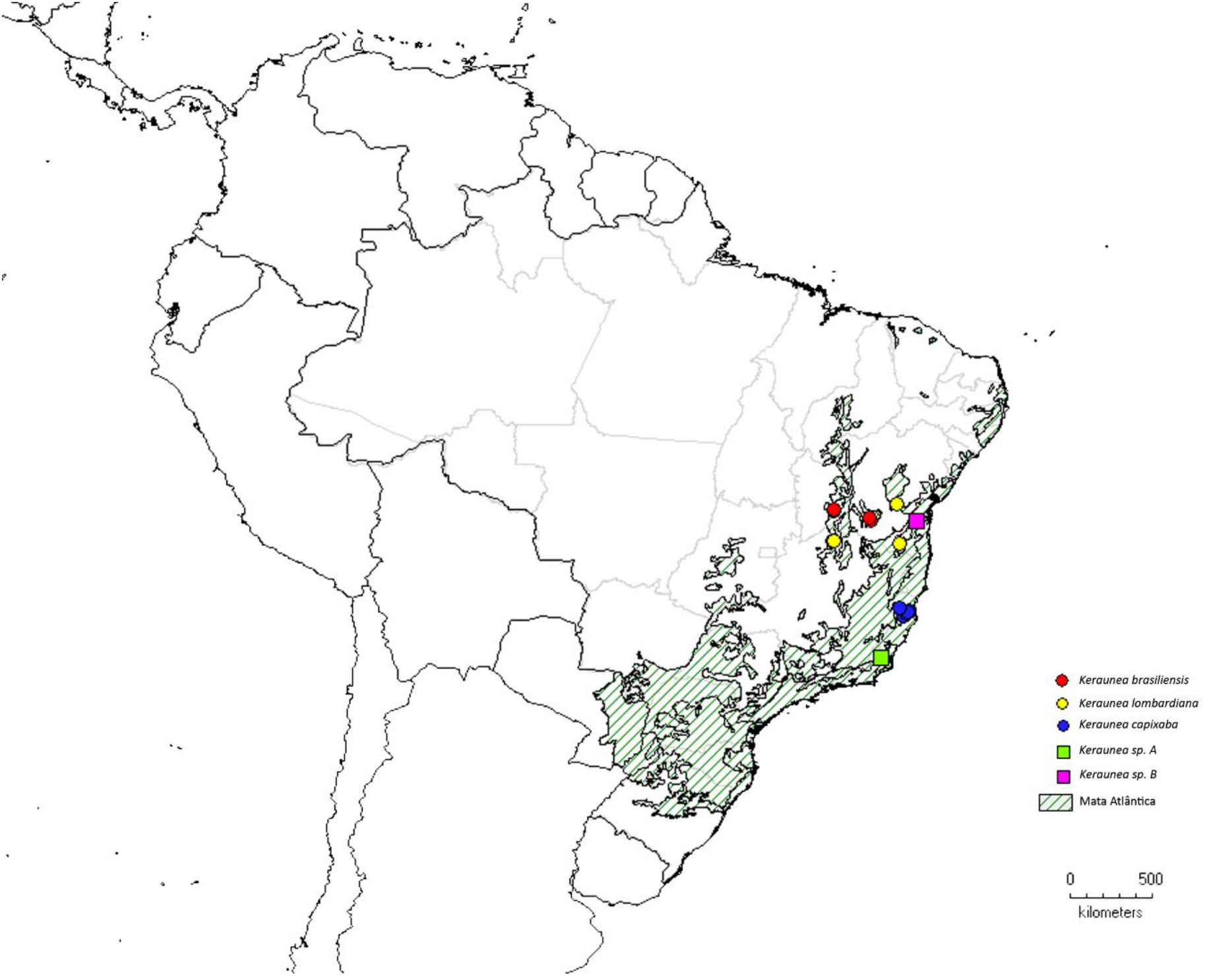
The distribution of taxa of *Keraunea* in relation to the Mata Atlântica biome, Brazil

*Keraunea capixaba* is definitely evergreen (Lombardi pers. obs.) and we postulate that the other moist forest species are also, but speculate that those from dry forest types *(K. brasiliensis* and *K. lombardiana*) may be deciduous in the dry season.

#### Scientific discovery of species

The earliest collection of *Keraunea* that we have traced was made nearly 100 years before the first species was named. This was *Rose & Russell* 19179 (NY) collected in Bahia in June 1915 and identified as *Keraunea* in 2017 by J. Kallunki. The second earliest collection known to us is *Santos* 2139 (CEPEC), also from Bahia (*K. sp*. B), followed by *Mori et al*. 11534 (NY), (*Keraunea lombardiana*). The type specimens of both *K. brasiliensis* and *K. lombardiana* were collected in 1997. The first 20 years of the 21^st^ century produced double the number of specimens collected in the entire 20th century, including *Keraunea* sp. A and all known specimens of *K. capixaba*. Since specimens of *Keraunea* have usually been identified and placed in families such as Solanaceae, Nyctaginaceae and Dichapetalaceae, where they have remained unidentified, it is likely that additional specimens of the genus exist in the herbaria of Brazil and elsewhere.

#### Key to the species of *Keraunea*

1. Primary stems (bearing flowering and fruiting spur shoots) drying bright white-yellow, surface exfoliating and rough…………………………….***K. brasiliensis***
2. Primary stems (bearing flowering and fruiting spur shoots) drying black, smooth, with or without very fine longitudinal lines… …………………………………………..2
3. Primary stems densely and conspicuously patent white hairy, hairs patent; distal leaves of the spur shoots narrowly elliptic, length: breadth ratio 3 – 4: 1, conspicuously densely white appressed hairy on both surfaces. Rio de Janeiro **sp. ……………………………………………………………………**….**A**
4. Primary stems glabrous or inconspicuously dark hairy; distal leaves of the spur shoots oblanceolate to broadly ovate, length: breadth ratio <3: 1, glabrous or if conspicuously hairy (*K. lombardiana*) only on upper surface. Bahia, Espirito Santo, Minas Gerais ……………………………………………………………………3
5. Distal leaves of spur shoots with adaxial surface scabrid, densely covered in conspicuous long, bright white, thick-based, stout hairs; leaves broadly ovate, 6.4 – 7.8 cm wide… ……………………………………………………***K. lombardiana***
6. Distal leaves of spur shoots with adaxial surface smooth, glabrous or sparsely covered in inconspicuous fine hairs; leaves elliptic, rhombic or oblanceolate, <6 cm wide…..4
7. Distal leaves of spur shoots with apex acuminate, secondary nerves 4 on each side of the midrib, stems and leaf margins hairy. Bahia ….…………….…………..***K*. sp. B**
8. Distal leaves of spur shoots with apex acute, secondary nerves 4 – 9 on each side of the midrib, leaves and stems glabrous. Espirito Santo & southern Rio de Janeiro………………………………………………..…………………..…***K. capixaba***

***Keraunea brasiliensis*** Cheek & Sim.-Bianch. Cheek & Simão-Bianchini (2013). Type: Brasil, Bahia, Mun. Caetité, Caminho da Fazenda Boa Vista para Urânio. 13° 59’ 35”S, 042° 12’ 27”W, alt. 560m. *Passos, Guedes, Stannard* and *Saar* in PCD 5263, fr. 8 Feb. 1997 (holotype SPF!; isotypes ALCB!, CEPEC!, K!). Fig. 4 & 7B, H&I.

*Scandent shrub* to c. 7 m tall, principal axes (bearing spur shoots) repeatedly branched, terete, solid, internodes 0.8 – 2.9 cm long, 2.2 – 2.5 mm diam., epidermis near shoot apex glossy yellow-brown, longitudinally wrinkled, inconspicuously and sparsely hairy, hairs fine, white, arched, 0.08 – 0.2 mm long; epidermis soon splitting longitudinally, exfoliating, surface becoming brownish white, corky, irregularly ridged, rough, lenticels not seen. Leaves of principal stems not seen. *Spur stems* (brachyblasts spur axis 2.7 – 4 cm long, internodes 2 – 4, first internode 1.7 – 2.8 cm long, second and third internodes 0.1 – 0.9 cm long, 1 – 1.5 mm diam.; axillary buds brown, subglobose, c. 1 mm diam.; indumentum dimorphic, 1) hairs fine, 0.12 – 0.25 mm long, arched-appressed, dense, forming a layer 0.1 – 0.2 mm deep, 2) sparser, erect, translucent, uniseriate hairs (0.12 –)0.2 – 0.25(– 0.3) mm long, the apex truncate, often red. *Distal leaves* drying pale green-brown, elliptic, rarely oblanceolate-obovate, 5.3 – 8.8 × 2.7 – 4.4 cm, apex and base acute, margin entire, secondary nerves 3 – 4(– 6) on each side of the midrib, on abaxial surface bright yellow-white, raised, ascending at c. 45 ° from the midrib +-straight, then bifurcating before attenuating, sometimes looping; tertiary and quaternary nerves inconspicuous, visible under magnification only, black, reticulate; on adaxial surface secondary nerves faintly visible, yellow-white; indumentum of adaxial surface with hairs colourless, not thickened, straight, sparse, 0.2 – 0.3 mm long, sparse, c. 0.5—1 mm^2^; abaxial surface with hairs as adaxial surface but 0.1 – 0.2 mm long, on nerves only. *Proximal leaves*, as the distal, but slightly smaller, elliptic-oblong, 4.8 – 5.7 × 1.9 – 2.3 cm, apex rounded, base acute. *Petioles* terete, 0.4 – 0.7 × 0.1 cm, indumentum as spur-shoots. *Inflorescences and flowers* not seen. *Infructescence* 1 – 3-fruited, peduncle 10 – 14 mm long, rhachis 0 – 2 mm long, rhachis internodes 0.2 – 1 mm long; free part of the pedicels 3 – 5(– 6) mm long. *Fruit* inserted on the accrescent bract; bract membranous, translucent, matt, elliptic, 54 – 62 x (2.5 –)2.8 – 3.2 mm, base and apex acute, indumentum inconspicuous, with a few translucent, fine, simple hairs 0.1 – 0.2 mm long scattered on the margins and on the proximal part, on and near the adnate pedicel; calyx persistent, sepals green in fruit, narrowly triangular 9.5 – 13 × 3 – 4 mm, the proximal united part cupular c. 5 mm diam., 1.5 mm deep, inner surface glabrous, outer surface moderately densely hairy, hairs white, fine, simple, slightly curved, 0.2 – 0.35(– 0.45) mm long; fruit green, inserted about the centre of the adaxial surface of the bract, mainly concealed within the calyx, subglobose 5 – 7 × 5 – 6 mm, epicarp glabrous, subglossy, thinly leathery, surface reticulate when dried due to endocarp projections; style persistent, single, 5 – 6 × 0.25 mm, apex bifurcate, T-shaped, branches 0.4 mm long; stigmas truncate; mesocarp spongiose with branching vascular bundles and some pulp between the endocarp (two pyrenes) projections; endocarps two; pyrenes plano-convex in outline, c. 11.5 × 6 × 2.5 mm, apices with two forward directed stout spines, bases rounded, closely appressed to each other ventrally, ventral surfaces flat; dorsally convex, each with a deep longitudinal groove, c. 5 lateral spines on each side, the remaining dorsal surface with longitudinal wings and other projections (Fig 4), each pyrene with two, single-seeded cells. *Seeds* arranged longitudinally, white, slightly curved, narrowing gradually from the rounded base to the acute apex (5.5 × 1.5 × 1.5 mm), cotyledons enveloped in endosperm, flat, appressed to each other, filling c. 50 % of the seed, radical apical.

### DISTRIBUTION

Brazil, Bahia.

### SPECIMENS EXAMINED

**BRAZIL. Bahia (BA)**, Mun. Caetité, Caminho da Fazenda Boa Vista para Urânio. 13° 59’ 35” S, 042° 12’ 27” W, alt. 560m. *Passos, Guedes, Stannard* and *Saar* in PCD 5263, fr. 8 Feb. 1997 (holo. SPF; iso. ALCB, CEPEC00077827 (3 sheets), K000979156); ibid. Caetité, Distrito de Maniaçú, estrada para São Timóteo, km 6, 22 May 2004, *Pereira-Silva* 9111 (HUEFS091115 (2 sheets) n.v.); ibid, Coribe, c. 5 km S em estrada de terra que cruza pequeno ramal que sai a 5.1 km E de Ponto d’água, a 24.4 km S de São Félix do Coribe na estrada para Coribe, fr. 10 April 2007, *Queiroz* 12707 (HUEFS118204, CEN00113310 n.v., SP444535 n.v.); Santa Maria da Vitória, ca. 7.7 km S de Santa Maria da Vitória na estrada para Lagoinha. Extremidade setentrional da Serra do Ramalho, fr. 13 Feb. 2000, *Queiroz* 5972 (CEPEC00113512 (2 sheets), HUEFS43719, SP4444534 n.v.)

### HABITAT

Caatinga, seasonal forest; 500 – 600 m alt.

### CONSERVATION STATUS

Known from four sites, with threats, *Keraunea brasiliensis* is here provisionally assessed as Endangered, EN B2ab(iii) according to the criteria of IUCN (2001). Area of Occupancy is estimated as 16 km² using 4 km² cells as advised by IUCN.

### PHENOLOGY

Fruiting in February and April-May; flowering unknown.

### ETYMOLOGY

Meaning coming from Brazil.

### VERNACULAR NAMES & USES

None are known.

### NOTES

*Keraunea brasiliensis* is the type and first described species of the genus (Cheek & Simão-Bianchini 2013). In the protologue, specimens of a separate species, *K. lombardiana*,described below, were included within *Keraunea brasiliensis*. However, it was stated that owing to the geographical, morphological and ecological separation between the specimens from Minas Gerais and that from Bahia, it might be that two species could be involved, not one (Cheek & Simão-Bianchini 2013). The additional specimens that have since become available support this hypothesis. The two species are readily separable from each other as indicated in the diagnosis (recognition features) of *K. lombardiana*.

*Keraunea brasiliensis* differs from the other three species of the genus recognised in this paper in that the primary stems are highly branched in all specimens (in the specimens of the other species they are unbranched, apart from the spur-stems), and the texture of the leaves is membranous or thinly chartaceous (vs thickly chartaceous). It is described as a scandent shrub, suggesting that the stems may not twine as is indicated in other species of the genus, e.g. *K. capixaba*

***Keraunea capixaba*** Lombardi Type: Brazil, Espirito Santo, Jaguaré Perto da comunidade São Jorge de Pádua. Estrada sentido para Fátima, 18° 54’ 29” S, 40° 08’ 44.9” W, fl. 25 Sept. 2013, *Siqueira* 891 (holotype CVRD; isotypes HRCB, K n.v., SP). Fig. 1, 3 & 6.

**Fig. 6.**
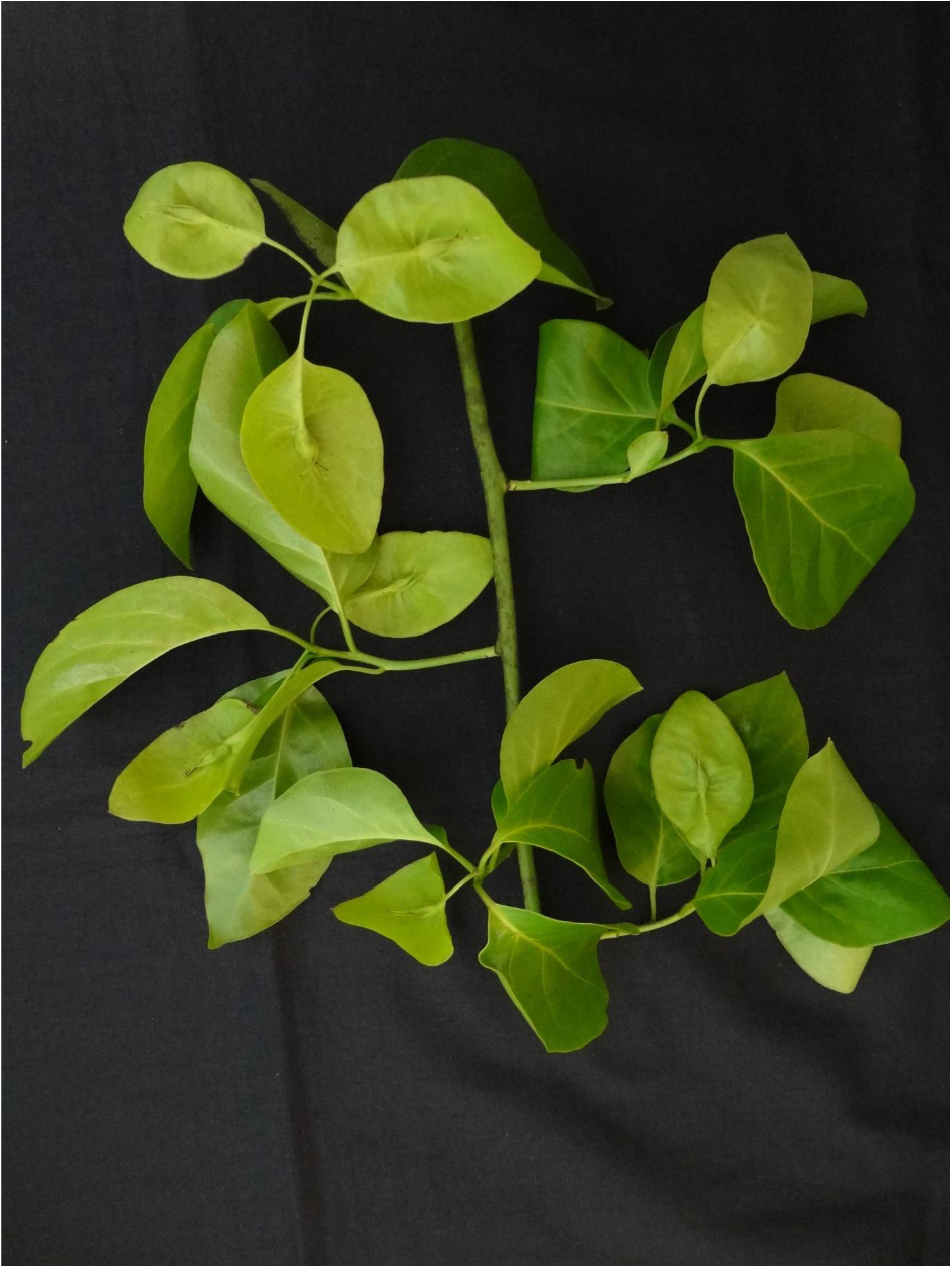
*Keraunea capixaba*. Section of primary stem with four fruiting spur-shoots, fruits immature. From *Siqueira* 893. Photo by Geovane Siqueira

### DISTRIBUTION

Brazil, Espirito Santo.

### SPECIMENS EXAMINED

**BRAZIL. Espírito Santo (ES)**, Jaguaré Perto da comunidade São Jorge de Pádua. Estrada sentido para Fatima, 18° 54’ 29” S, 40° 08’ 44.9” W, fl. 25 Sept. 2013, *Siqueira* 891 (holotype CVRD14565; isotypes HRCB76194, HRCB76196, RB608435, RB00895208, K n.v., SP476897); ibid 18° 54’ 29” S, 40° 08’ 44.9” W, fr. 16 Oct. 2013, *Siqueira* 893 (CVRD14570; ESA137022, HRCB76199, K001275507, MBML048106, MO101180015, NY02687786, RB00895202; SPF234754); ibid Jaguaré Perto da comunidade São Jorge de Pádua. Estrada sentido para Fatima, 18° 54’ 29” S, 40° 08’ 44.9” W, fl. 25 Sept. 2013, *Folli* 7117 (CVRD14563, HRCB 76194, MO101180011, NY02687787, RB00895205, K n.v.); ibid Sooretama, Rodovia, Distro de Bom Jardin, fl. fr. 1 Sept. 2012, *Assis & Freitas* 3340 (HRCB76193, MBML044646 (2 sheets); ibid, Rio Bananal, Alto Bananal, propr.: Jonas Graci, 19° 14’ 56”S, 040° 24’ 59”W, fr. 25 April 2007, *V. Demuner; T. Cruz, R. R. Vervloet, M. Belisario & Elias Bausen*, 3799 (BHCB124910 n.v., MBML30094 n.v.); Sooretama, Reserva Natural Vale, Estrada Barro Roxo a Córrego Rodrigues, fl. fr. 8 Oct. 2014, *Folli* 7273 (CVRD15122 n.v., RB1103439); ibid, Reserva Biológica de Sooretama. Mata de Tabuleiro, 19 May 2015, *Covre* s.n. (SAMES03696); ibid, Nova Venécia, Fazenda Santa Rita. Ao pé da pedra da torre, fr. 2 Feb. 2018, *Gurtler & Dutra* in JAC 371 (VIES036853 2 sheets).

### HABITAT

Borders of secondary or disturbed seasonal forests (mata de tabuleiro) in the Atlantic Coastal Forest, on coastal plains at c. 100 m alt. (Lombardi 2014).

### CONSERVATION STATUS

Endangered with fewer than 250 individuals and an extent of occurrence estimated to be less than 500 km^2^ (Lombardi 2014).

### PHENOLOGY

Flowering in September.

### ETYMOLOGY

Meaning (Brazilian, indigenous origin) for people born or things native to the state of Espírito Santo.

### VERNACULAR NAMES & USES

Canema Branca; uses if any unknown

### NOTES

Since the protologue was published (Lombardi 2014), five specimens additional to the four originally known have come to light, extending the range slightly. With nine specimens, this species of the genus is the most frequently collected.

*Keraunea capixaba* is remarkable for being almost entirely glabrous, with hairs only on the sepal margins (Lombardi 2014). The fruits are unusual in not having a reticulated, but a smooth surface (Cheek pers. obs. 2023) The flowers are reported to be highly perfumed (*Assis & Freitas* 3340, *Folli* 7117) and are numerous and large (Fig. 3). The species is reported to be a stem twiner in field collection notes (“trepadeira” and “liana volúvel”).

The isotype cited at K in the protologue has not been found, but the paratype (*Siqueira* 893, K) was the basis for DNA studies including this species.

Data on this species in cultivation is given under the generic notes above.

***Keraunea lombardiana*** Cheek sp nov. Type: Brazil, Minas Gerais, Januária, Distrito de Fabião, junto ao abrigo do Malhador 15° 07’ 85” S, 44° 15’ 17” W. fr. 25 May 1997, *Lombardi* 1819 (holotype BHCB 37301; isotype K000593363). Fig. 5 & 7A, C-G.

*Keraunea brasiliensis* pro parte quoad *Lombardi* 1819 & *Lombardi* 2107 (Cheek & Simão-Bianchini 2013)

*Climber to 4 m tall over low trees. Primary stems* climbing, terete, fistular, dark purple, finely longitudinally ridged, ridges rounded, c. 15 – 20, internodes 5 – 7.5 cm long, 4 – 5 mm diam., cicatrices (leaf scars) narrowly heart-shaped, c. 2.5 × 2 mm, axillary bud rounded, with scales, c. 1 × 1.8 mm, minutely and inconspicuously hairy; hairs 20 per mm^2^, translucent, dimorphic, 1) narrowly tapered, unicelled hairs 0.25 – 0.4 mm long, interspersed with 2) truncate, stout, multicellular hairs 0.1 – 0.15 mm long. *Leaves* of primary stems ovate to lanceolate, 9.5 – 11 × 5.5 – 7 cm, apex acuminate-acute, base truncate or very shallowly cordate, lateral nerves 5 on each side of the midrib, arising straight at 50 – 70 ° from the midrib, branching about half way to the margin and uniting with the secondary nerves above and below, forming a looping inframarginal nerve; tertiary nerves reticulating in the marginal part of the blade; adaxial indumentum of stout, acute, erect, white hairs 0.6 – 1 mm long, 0.05–0.07 mm wide, arising from a conical base c. 0.1 × 0.1 mm, density 4 – 6 mm^2^; abaxial indumentum as adaxial but hairs (0.08 –)0.5 mm long, bases reduced. Petioles articulated at base, canaliculate (0.9 –)2 × 0.15 – 0.2 cm, the base abruptly dilated, cordate, 3 mm wide, indumentum as stem. *Spur stems* (brachyblasts) arising c. 2 mm above the axillary buds, subtended by a whorl of 4 – 6 scale-leaves; scale leaves moderately caducous, becoming larger distally, +-rhombic (the most distal +-orbicular), 3.5 – 5 × 3.5 – 5 mm, broadest in the distal half, apex obtuse or acute, marginal band black, c.0.5 mm wide, inner surface glabrous, drying green, outer surface and margin with appressed, bright white, crinkled hairs c. 0.25 mm long, spur axis 2.7 – 4 cm long, internodes 2 – 3, first internode 2 – 2.2 cm, second and third internodes 0.2 – 0.9 cm long, 1.7 – 2 mm diam.; axillary buds brown, subglobose, 1.5 – 3 × 1.5; indumentum with hairs fine, white, 0.25 – 1 mm long, arched, forming a dense blanket, mixed with a few sparse and inconspicuous, erect translucent multiseriate hairs 0.2 – 0.25 mm long. *Distal leaves* drying black-grey, thickly chartaceous, broadly ovate, 8.5 – 10.5 × 6.4 – 7.8 cm, apex rounded to obtuse, base broadly obtuse, rounded, truncate or shallowly cordate, margin appearing minutely toothed and scabrid due to hair bases; secondary nerves (5 –)6 – 7 on each side of the midrib, on abaxial surface bright yellow-white, strongly raised, ascending at c. 45° from the midrib +-straight in the proximal half of their length, gradually arching upwards, becoming parallel with the margin, uniting via tertiary nerves with the secondary nerves above, before attenuating; tertiary nerves conspicuous, subscalariform, uniting the secondary nerves; quaternary nerves reticulate, forming areolae 2 – 4 mm diam., indumentum as the primary stem leaves, but adaxial surface with hairs arching, appressed, 1.2 – 1.3 mm long, bases 1.12 mm diam., density 3 – 4(– 5) mm^2^, with age breaking above the base, and becoming scabrid. *Proximal leaves* as dorsal but elliptic, 3 – 6 × 2 – 3.7 cm, apex and base rounded, petiole 0.6 – 0.9 cm long. *Inflorescence* (immature) terminal on short leafy, spur shoots, corymbose, 7 – 8 mm, long, 3 – 5-flowered, indumentum as petiole; peduncle 1 – 1.2 mm; pedicel 2 – 3 mm long; bract inserted at apex of pedicel, elliptic, 3.6 – 3.9 × 2 – 2.2 mm, base decurrent by 0.5 mm. Bracteoles absent or inconspicuous, calyx tube campanulate, 0.5 – 0.9 mm long; lobes five, narrowly triangular to narrowly lanceolate 2.8 – 3.8 × 0.6 – 0.9 mm, pinnately nerved, outer surface and margin with indumentum as leaf-blade, inner surface glabrous; corolla c 3.6 mm long; corolla tube c 0.5 mm long, corolla lobes oblong-elliptic 3.5 – 4 × 1.5 – 1.7 mm, apex rounded, with a few simple hairs 0.25 – 0.5 mm long near the apex, midpetaline only weakly developed as a darker area, glabrous; androecium with five stamens, epipetalous, inserted at base of corolla tube, alternating with petals, glabrous; filaments 0.3 – 0.4 mm long, anthers basifixed, cells 2.5 – 2.6 × 0.7 mm, introrse, connective distal, 0.4 – 0.6 mm long, apex globose; disc 0.1 mm thick; gynoecium superior 1.8 × 0.7 – 0.8 mm, glabrous, ovary subglobose, style single, conduplicate, unbranched, stigmas two, truncate; ovary 2-locular, locules biovulate. *Infructescence* 1 – 3-fruited, peduncle 4 – 6 mm long, rhachis 0 – 6 mm long, rhachis internodes 0.5 – 1 mm long, pedicel 5 – 6(– 10) mm long, indumentum as spur stem. *Fruit* inserted near the centre of the accrescent bract; bract papery, shining, bullate, elliptic, (37 –)43 – 55 × 24 – 33 mm, indumentum on both surfaces as spur stem, hairs fine +-erect, 0.3 – 0.6 mm long; calyx accrescent, green, united at base forming a shallow cup 5 mm diam., lobes 8 – 10 mm long, proximal 4 mm c. 2 mm wide, distal portion 1 mm wide, tapering to acute apex, abaxial surface with conspicuous, broad, raised, yellow midrib, densely white hairy, hairs 0.25 – 0.1 mm long, conspicuous at margins; adaxially glabrous. *Fruit* green in life, drying brown, inserted about the centre of the adaxial surface of the bract, subglobose, 8 10 × 8 – 10 mm, surface with irregular, mainly longitudinal ridges, glabrous, style persistent, single, 3 – 6 mm long. Mesocarp spongiose with vascular branches and sparse pulp; endocarps 2, woody, plano-convex, c. 9 × 9 × 5 mm, ventral faces flat, closely appressed, the circumference with a wide, translucent wing supported by short transverse ridges; dorsal faces with blade-like wings, locules two, each with a single seed, Seeds c. 4 × 1.8 × 1.8 mm.

### RECOGNITION (Diagnosis)

Similar to *Keraunea brasiliensis* Cheek & Sim.Bianch., distinct in the leaves broadly ovate, thickly papery, 6.4 – 7.8 cm wide (vs elliptic or obovate-oblanceolate membranous or thinly papery, 2.7 – 4.4 cm wide), the primary axes (bearing spur-shoots) fistular, dark purple, smooth (vs solid, bright yellowish white, 2.2 – 2.5 mm diam., rough and exfoliating), fruits 8 – 10 mm diam. (vs 5 – 7 × 5 – 6 mm diam.), mericarps with a circumferential wing, lacking spines (vs mericarps spiny, lacking wings).

### DISTRIBUTION

Brazil, northern Minas Gerais and southern Bahia.

### SPECIMENS EXAMINED

**BRAZIL. Minas Gerais (MG)**, Januária, Distrito de Fabião, junto ao abrigo do Malhador 15° 07’ 85” S, 44° 15’ 17” W. fr. 25 May 1997, *Lombardi* 1819 (BHCB, K00139055); ibid, junto Abrigo do Malhador entre 15° 07’ 16” - 15° 08’ 57” S, 44° 15’ 20” - 44° 14’ 13” W, fl. buds 26 Oct. 1997, *Lombardi* 2107 (BHCB, K000593363).

**Bahia (BA)**, Itambé, Rod. BA-265, trecho Itapetinga/Caatiba. Faz. Serra Verde, a 17 km da Rod. BR-415, 15° 15’ 33.58” S, 40° 37’ 31.39” W, fr. 14 March 1979, *Mori, dos Santos, Thompson* 11534 (CEPEC00016207 n.v., MO0100961141 n.v., NY03147645, RB00725192); Vicinity of Machado Portello, 13 ° 8’ 25” S, 40 ° 46’ 10” W, fr. 19-23 June 1915, *Rose & Russell* 19979 (NY01092952)

### HABITAT

Dry or mesophilic forest, often disturbed, sometimes on stony substrate.

### CONSERVATION ASSESSMENT

Known from three locations, each with threats and thought to be unprotected, therefore it is provisionally assessed here as Endangered. In the type location, in Minas Gerais, JAL observes that at the time of collections he made in 1997, the habitat in which the specimens were found was secondary, and close to roads. Since that time new protected areas have been created in the area, but these do not include the known sites for *K. lombardiana*. However, it is possible, even likely, that the species may yet be found in those protected areas.

### PHENOLOGY

Flower buds in late October, fruiting the following May & June

### ETYMOLOGY

Named in honour of Julio A. Lombardi (1961-) plant systematist and taxonomic specialist in Neotropical Celastraceae, Vitaceae, Oleaceae, and other families. He is professor at Universidade Estadual Paulista (UNESP), Campus de Rio Claro, Instituto de Biociências. He collected two of the four known specimens of the species named for him, including the type specimen.

### VERNACULAR NAMES & USES

None are known.

### NOTES

This species was formerly included within *Keraunea brasiliensis* (Cheek & Simão-Bianchini 2013)). However, the protologue highlighted the geographical and morphological separation between the type, in Bahia and the southern elements, in Minas Gerais, and it was suggested that with more data, it might be possible to evidence that two species were present, as is shown here.

*Keraunea lombardiana*, of all the known species of the genus, has indumentum that most resembles that usual in Boraginales, with stout-based, white, bristly hairs, which break above the base, leaving a cylindric shaft, making the leaves scabrid on the adaxial surface. The conical bases of the hairs give the leaf margin the appearance of being finely dentate. only flowers recorded to date are highly immature (Fig. 7D-F).

**Fig. 7.**
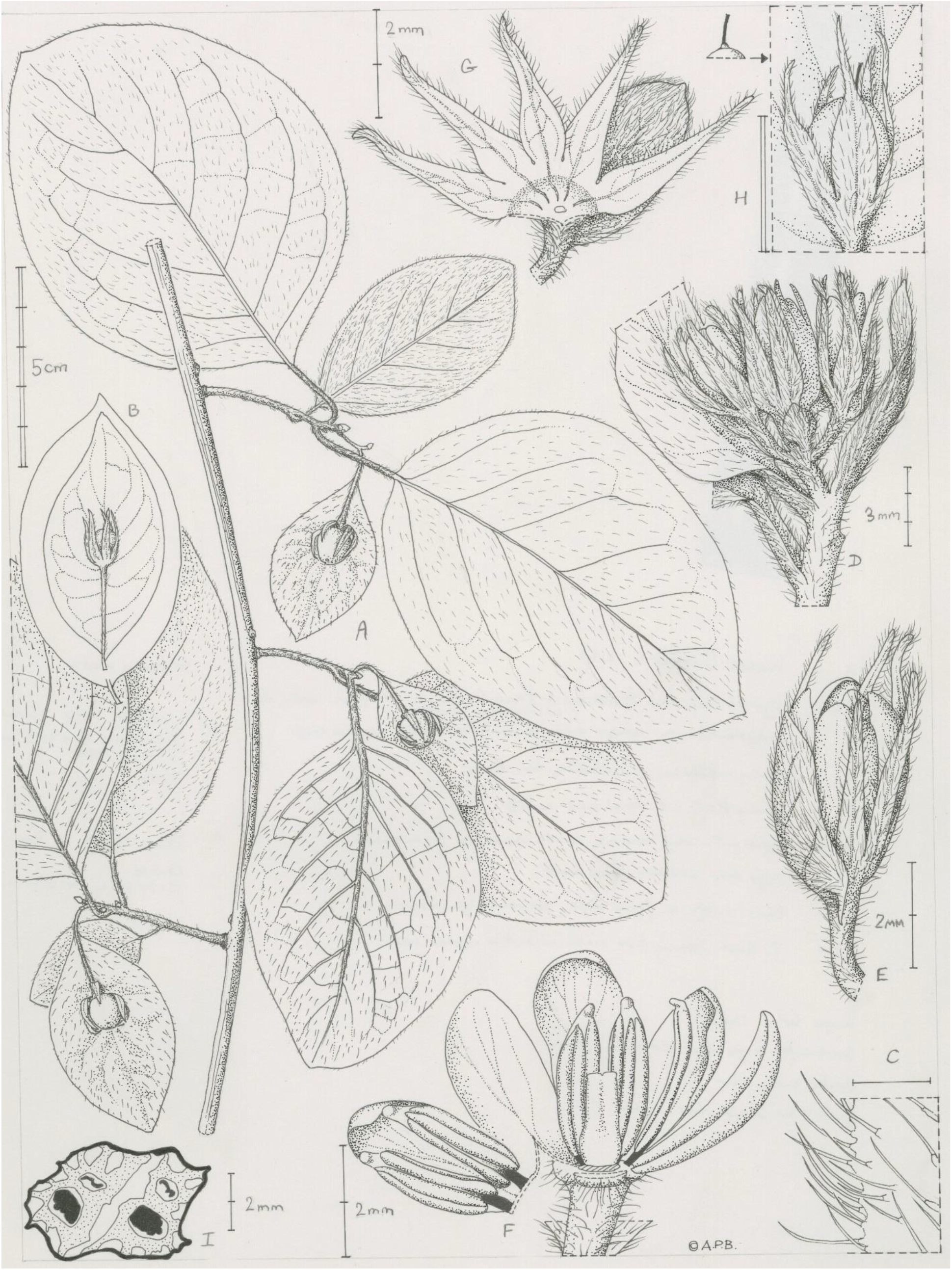
*Keraunea lombardiana* A. Habit; C. Hairs of young leaf-adaxial surface and margin; D. Inflorescence, before anthesis; E. Flower (pre-anthesis) with bract. F. Flower (pre-anthesis) dissected, calyx removed; G. Calyx (dissected), receptacle and bract (pre-anthetic flower). *Keraunea brasiliensis* B. fruit with outline of typical leaf for comparison with *K. lombardiana*; H. Fruit (immature) on bract; I. Fruit, transverse section, showing two winged, woody pyrenes (dotted) each with one functional and one aborted seed. A, C-G from *Lombardi* 1819 (*Keraunea lombardiana* K); B, H & I from *Passos et al*. 5263 (*Keraunea brasiliensis*), K; scale bars: A & B= 5 cm; C = 1 mm; D= 3 mm; E, G, F & I = 2 mm; H = 2cm. All drawn by Andrew Brown.

The broad, very thick, densely hairy, leaves with bases broadly obtuse to cordate help to distinguish this species from others of the genus, which all have finer, much more sparse and inconspicuous hairs on the leaves, or which are glabrous.

The fruits of *Keraunea lombardiana* are larger than those of *K. brasilensis* and differ in that their endocarps have an intact marginal wing (Fig.5), rather than the wings being divided, forming marginal spines or teeth. The endocarps of *K. brasilensis* differ from the first additionally in having a pair of long apical spines, and a deep longitudinal groove on the dorsal surface between the two seed cells (Fig. 4).

***Keraunea* sp. A**

### DISTRIBUTION

Brazil, Rio de Janeiro

### SPECIMENS EXAMINED

**BRAZIL. Rio de Janeiro (RJ)**, Cardoso Moreira, Moreira Santíssimo, em fragmento de mata, Fazenda Borges, fl. 10 Sept. 2013, *Costa* 257 (RB barcode 00852871).

### HABITAT

Forest fragment.

### CONSERVATION STATUS

Likely to be Critically Endangered since a single site is known on current evidence, and it appears to be unprotected.

### PHENOLOGY

Flowering in September.

### VERNACULAR NAMES & USES

None are known.

### NOTES

This is a particularly distinct species which is known to us only from a digital image from which we are unable to prepare a complete description. The narrowly elliptic leaves, covered on both surfaces by long, conspicuous, seeming non-hispid white hairs are seen in no other species. The species is also unique in that the reduced leaves which in other species are restricted to the base of the spur shoots as bud-scales, appear to be scattered along the proximal part of the spur shoot, and foliose (albeit reduced in size), below the fully-formed distal leaves.

This is the southernmost taxon of the genus known to us at present.

***Keraunea* sp. B DISTRIBUTION**. Brazil, Bahia.

### SPECIMENS EXAMINED

**BRAZIL, Bahia (BA)**, Ba. Rod. Ipiaú-Ibirataia, trepadeira fl. branca. Plantação de cacau, fl. 13 Nov. 1971, *T*.*S. Santos* 2139 (CEPEC8062).

### HABITAT

Plantation of *Theobroma* (cacao). Presumably this site had the remains of an evergreen forest fragment.

### CONSERVATION STATUS

This taxon is likely to be Critically Endangered since currently only a single site is known, which is unprotected as far as biodiversity conservation is concerned.

### PHENOLOGY

Flowering in November.

### VERNACULAR NAMES & USES

None are known.

### NOTES

This species has the hairy, purple black, smooth primary stems seen in *Keraunea lombardiana* (in *K. capixaba* the stems are glabrous). They differ in that the distal spur leaves of *K. sp. B* lack the conspicuous, thick based white hispid hairs seen on the adaxial surface of *K. lombardiana* (hairs can be seen in *K*. sp. B, but they appear to be finer, dark and at the leaf margins). The two taxa may be closely related. It is even possible that the differences seen are purely stage-related since the spur-stem leaves of *K*. sp. B are all young and freshly formed (the specimen is in the flowering state), while those of the specimens available of *K. lombardiana* either have immature flowers (the spur leaves not yet developed) or have fruits (with spur leaves months older than in *K*. sp. B). With only a digital image and without direct access to the specimen it is not possible to be certain of the status of this plant.

## Discussion

### *Keraunea’s* taxonomic affinities in Ehretiaceae

According to the molecular analyses, *Keraunea* is reconstructed as a highly supported clade within Ehretiaceae, disputing the previous results from Muñoz-Rodriguez *et al*. (2022), which reconstructed it as two lineages, one in Ehretiaceae and another in Malpighiaceae. Our BLAST results corroborate the scenario presented by Almeida et al (submitted), where the misplacement of *K. brasiliensis* in Malpighiaceae was possibly due to a mix of leaves fragments in the type specimen, since new sequences from the same specimen (Passos PCD 5263) matched those of Ehretiaceae in all markers. Not surprisingly, the discovery that *Keraunea* should be placed outside the Convolvulaceae (Simões *et al*. 2022) explains the former difficulties placing the genus within the tribal classification of that family (Cheek & Simão-Bianchini 2013). Placement within Ehretiaceae (Boraginales) has is also confirmed by morphological evidence. Initially, this seemed surprising, since the climbing habit and presence of a bract is not included in the morphological circumscription of the family (Luebert *et al*. 2016). However, in the more extensive treatment of Ehretiaceae by Gottschling *et al*. (2016) (albeit with the inclusion of Cordiaceae and Hoplestigmataceae), it is reported that some species of *Rochefortia* Sw. and *Bourreria* P. Browne are also scandent or lianescent, and a bract is reported in *Cortesia* (Mirra *et al*. 2022), here resolved as one of the possible sister genera to *Keraunea* (Fig. 2). The inflorescence and flower structure of *Keraunea* fit the existing circumscription of Ehretiaceae. The presence of a hair complement of both unicellular and uniseriate, sometimes gland-tipped trichomes (so far reported only in one species of the genus, *Keraunea brasiliensis*, see below) is also consistent with Ehretiaceae (Gottschling *et al*. 2016). The robust hairs with a raised multicellular base, sometimes with cystoliths in accessory cells, which produce a scabrid surface when the hairs are fractured with age (see *Keraunea lombardiana* account below) are indicative of Boraginales as a whole. However, *Keraunea capixaba* is almost entirely glabrous (Lombardi 2014).

At generic level, the majority of the lineages (genera) of pantropical Ehretiaceae are, like *Keraunea*, Neotropical, all being endemic to or with their major centre of species diversity in, S America, except for *Halgania* which is Australian. *Bourreria, Cordia* and *Ehretia* all show species radiations in the Old World.

Wiegend *et al*. (2014) report that the presence of 4 pyrenes (known as mericarps when they become physically divided from each other in the fruit as separate units) is ancestral (plesiomorphic) in the Ehretiaceae. Fusion of mericarps, or syn-mericarpy has evolved independently more than once in Ehretiaceae (Gottschling *et al*. 2014). Either all four pyrenes fuse together into one unit, or as we show in this paper occurs in *Keraunea*, two pairs of pyrenes fuse to form two 2-seeded pyrenes (or endocarps). These “two parted endocarps” occur in one subclade within *Ehretia* and apparently independently, in the genus *Cortesia*. (Gottschling *et al*. 2014). *Cortesia* is sister to the clade that includes *Keraunea* and *Halgania*, and these, then, form a clade sister to *Ehretia*.

These relationships are morphologically supported by the presence, uniquely within Ehretiaceae (in which bracts are otherwise absent (Gottschling *et al*. 2016)), of an involucral bract in *Cortesia*. Both the two parted endocarps and the presence of an involucral bract are flagged as apomorphies for *Cortesia* (Gottschling *et al*. (2014: Fig. 2)). Considering the molecular phylogenetic analyses, we hypothesise that the two parted endocarps and bracts are, in this context, homologous, representing synapomorphies for the clade that includes the genera *Cortesia, Halgania* and *Keraunea* (we conjecture that the bracts are later lost in *Halgania*).

Despite this morphological support for a sister relationship of *Cortesia* and *Keraunea*, the two genera are otherwise dissimilar in leaves, habit and habitat. *Cortesia* is a halophytic shrub of Argentina with cuneate, tridentate leaves, spatulate sepals and white drupes (Mirra *et al*. 2022). There is certainly no morphological basis for uniting the two genera. The phylogenetic relationship among these two genera remains unclear, being reconstructed either as sister to each other, or with *Keraunea* closer to *Halgania*, then *Cortesia*. This uncertainty is possibly caused by the low amount of data for *Cortesia*, with fragments of sequences for only two (ITS and *trn*L-F) of the four markers used here. To improve the resolution among these genera, a more extensive molecular sampling for *Cortesia* is required, ideally based on nuclear data (e.g., following Baker *et al*. 2021).

Despite the strong similarity of *Keraunea* to *Neuropeltis* (Convolvulaceae) in the fruit arrangement and other features, the evidence shows that these are due entirely to convergence and not shared evolutionary history. This is an interesting discovery that opens new avenues for re-evaluating the current knowledge on character evolution and systematic relationships within Convolvulaceae, Ehretiaceae and, in fact, Angiosperms as a whole. Especially in cases where family and generic placements have been proposed with incomplete morphological data or poor molecular phylogenetic sampling, it is important to keep open the possibility to revisit these hypotheses in the future, and assume that the taxon can later be transferred to a different group based on newly available evidence. This is, for instance, the recent case of *Distimake vitifolius* (Burm. f.) Pisuttimarn & Petrongari, in Convolvulaceae (Pisuttimarn *et al*. in press), where incomplete molecular data and the assumption of six-zono-colpate pollen being synapomorphic of the genus *Camonea* Raf., led to an erroneous placement of the species in that genus (Simões & Staples 2017).

Therefore, while taxonomists are often faced with the difficult task of having to make decisions on the (scarce) evidence at hand at a particular moment, as more evidence becomes available, a reinterpretation of the data is always possible, and sometimes necessary, to resolve the placement of curiously ambiguous taxa, as is the case of *Keraunea*, where preliminary molecular data suggested first its incorrect placement in Convolvulaceae (Simões *et al*. 2022), then its placement in Malpighiaceae and Ehretiaceae (Muñoz-Rodriguez *et al*. 2022), and now it is finally confirmed to belong in Ehretiaceae with both morphological and molecular support. The material now named *Keraunea* had been known to several scientists in Brazil for some years before it was published, including by one of the authors of this paper, but had not been described because there had been no conclusion on which family to place it in. However, it is valuable to formally name new taxa, even when their final placement in a classification is uncertain due to lack of sufficient molecular phylogenetic data (e.g. Darbyshire *et al*. (2023). This is because, until a taxon has a scientific name, it is essentially invisible to science (Cheek *et al*. 2020). Since it was discovered (Cheek & Simão-Bianchini 2013), *Keraunea* has become visible to science, being widely referred to (see Introduction), and as it was brought to the attention of the taxonomic community, it thus facilitated the discovery of more species (Lombardi, 2014).

## Conclusions

*Keraunea* has being challenging botanists for more than 100 years, since its first collection in 1915, to its formal description in Convolvulaceae in 2013 (Cheek & Simão-Bianchini, 2013) to the recent tentative placement in Malpighiaceae (Muñoz-Rodriguez et al, 2022). Our results now shed light on the evolutionary history of this genus, unequivocally placing *Keraunea* within Ehretiaceae and confirming it as a monophyletic lineage. We also present a new species, *K. lombardiana*, distinctive by broadly ovate leaves, fistular and dark purple bearing spur-shoots, and winged (not spiny) mericarps. Despite the significant improvements to the natural history of *Keraunea* presented here, in many respects, this genus remains incompletely known. Two of the five species known in the genus lack formal descriptions and not one of the named species has both anthetic flowers and fruits described. Studies on pollen, germination, anatomy, embryology and phytochemisty are entirely lacking. Field observations of pollination, seed dispersal and phenology are also required. However, the most urgent requirement is undoubtedly a formal taxonomic revision with full IUCN (2012) extinction risk assessments for each taxon. This is urgent because all the taxa that we present in this paper (and more can be expected to be found) appear to be highly range restricted and generally not to occur in protected areas, and so are likely to be highly threatened. We hope that future urgent assessments might facilitate the incorporation of these species in protected areas and the development of individual or area-based species conservation action plans (e.g. Couch *et al*. 2022) to reduce the risk of their extinction.

## Acknowledgements

Leo-Paul Dagallier is thanked for providing photos from NY. We thank Eimear Nic Lughadha for reviewing an early draft of the ms. We also thank Lazslo Csiba for lab support

Two anonymous reviewers are thanked for comments on an earlier version of the manuscript.

The authors declare that they have no conflict of interest.

## Notes

### Competing Interest Statement

The authors have declared no competing interest.

